# Link between the numbers of particles and variants founding new HIV-1 infections depends on the timing of transmission

**DOI:** 10.1101/404913

**Authors:** Robin N. Thompson, Chris Wymant, Rebecca A. Spriggs, Jayna Raghwani, Christophe Fraser, Katrina A. Lythgoe

## Abstract

Understanding which HIV-1 variants are most likely to be transmitted is important for vaccine design and predicting virus evolution. Since most infections are founded by single variants, it has been suggested that selection at transmission has a key role in governing which variants are transmitted. We show that the composition of the viral population within the donor at the time of transmission is also important. To support this argument, we developed a probabilistic model describing HIV-1 transmission in an untreated population, and parameterised the model using both within-host next generation sequencing data and population-level epidemiological data on heterosexual transmission. The most basic HIV-1 transmission models cannot explain simultaneously the low probability of transmission and the non-negligible proportion of infections founded by multiple variants. In our model, transmission can only occur when environmental conditions are appropriate (e.g. abrasions are present in the genital tract of the potential recipient), allowing these observations to be reconciled. As well as reproducing features of transmission in real populations, our model demonstrates that, contrary to expectation, there is not a simple link between the number of viral variants and the number of viral particles founding each new infection. These quantities depend on the timing of transmission, and infections can be founded with small numbers of variants yet large numbers of particles. Including selection, or a bias towards early transmission (e.g. due to treatment) acts to enhance this conclusion. In addition, we find that infections initiated by multiple variants are most likely to have derived from donors with intermediate set-point viral loads, and not from individuals with high set-point viral loads as might be expected. We therefore emphasise the importance of considering viral diversity in donors, and the timings of transmissions, when trying to discern the complex factors governing single or multiple variant transmission.

## INTRODUCTION

Characterising the strong bottleneck that occurs during HIV-1 transmission, and understanding the role of selection in determining which viral variants are transmitted, are important for HIV-1 prevention strategies [1]. It is now well established that most infections are founded by one or few distinct viral variants (e.g. [2–8]), with each of these variants referred to as a transmitted/founder (T/F) virus. One T/F virus might naïvely be assumed to mean one T/F viral particle. However, it is currently unknown whether each T/F virus results from the successful transmission of a single viral particle, or multiple viral particles of the same variant, and as a corollary, how the number of viral particles founding an infection relates to the number of T/F variants. To avoid potential confusion, throughout we avoid using the term “virus”, and instead refer to viral particles or viral variants, as appropriate (see Glossary).

The observation that most HIV-1 infections are founded by only one or a few variants has been used as evidence for a strong selective bottleneck at the point of transmission, giving hope that signatures of transmission can be found and exploited when designing vaccines (e.g. [1,9,9]). However, the extent to which selection influences which viral variants are present at the start of an infection is a source of current debate [11–15]. It has also been observed that infections founded by multiple variants tend to have higher set-point viral loads (SPVLs) than those founded by single variants, with the possible implication that multi-variant transmission might be a trait associated with recipient individuals [16].

However, the hypotheses that a small number of T/F variants is indicative of selection, and that multi-variant transmission might be driven by recipient host factors, are missing explicit consideration of the complex interplay between viral load, viral diversity, and the timings of transmissions from infector individuals (donors) within a population. For a single donor at a fixed point during infection, the number of variants transmitted to a recipient is expected to be higher if a larger number of viral particles are transmitted. However, once the possibility of transmission occurring at any point during a donor’s course of infection is taken into account, it is not necessarily the case there is a simple link between these two quantities if multiple transmissions are considered. This is because the viral load typically varies by orders of magnitude during the course of an untreated infection, and viral diversity tends to increase as an infection progresses [17–19]. For example, early in an HIV-1 infection, the viral load is typically high but viral diversity is usually low [20], whereas during chronic infection the viral load is lower but diversity is typically higher. As a consequence, the relationship between the numbers of T/F variants and the numbers of T/F particles in a recipient population is likely to depend not only on selection and recipient host factors, but also on the compositions of variants in donors and the timings of transmissions.

Here we present a probabilistic model, informed by within-host deep-sequencing [18] and population-level [21] data, to investigate the likely relationship between the numbers of variants and the numbers of particles founding new sexually transmitted infections in untreated populations, as well as the link between donor SPVLs and the numbers of T/F variants among recipients. We also consider the impact that selection, and a bias towards early transmission (due to treatment and/or other behavioural factors), might have on the compositions of new infections.

Considering the timings of transmissions explicitly will make it easier to deduce the relative importance of selective and non-selective bottlenecks during transmission within different risk groups. The timings of transmissions might also provide an explanation for some perplexing results, such as the proportion of multi-variant transmissions in some studies of populations of men who have sex with men (MSM) being comparable to standard estimates in heterosexual populations [3,6–8] despite evidence for weaker selection during MSM transmission [8].

## RESULTS

### A probabilistic model of transmission

To characterise the relationship between the numbers of viral particles and viral variants that found infections in a population, we first developed a probabilistic model describing heterosexual transmission from a single untreated donor at a fixed time during infection, which we then scaled up to a population of untreated donors. Computing code for running our model can be accessed at https://github.com/robin-thompson/MultiplicityOfInfection.

A schematic of the probabilistic model for a single transmission event is shown in Figure 1. In brief, the expected numbers of particles and variants that are successfully transmitted depend on the viral load and the distribution of variants within the donor at the time of transmission. We account for the observations that HIV-1 is only transmitted rarely [22], but when transmission does occur, multiple viral variants found the new infection reasonably often [2,4,4]. These observations cannot be captured simultaneously by simple models, such as binomial transmission models, since in these models a low probability of transmission predicts that, when transmissions occur, they will only be with single particles and therefore single variants (see [4], and also Text S1 – Binomial Models of Transmission). To reconcile these two observations, we assume that transmission can only occur in a small fraction, *f*, of potential transmission acts, when environmental conditions are appropriate. This is supported by observations that HIV-1 is most likely to be transmitted when a potential recipient is experiencing abrasions in the genital tract, genital inflammation, or coinfection with another pathogen [12,23–27].

**Figure 1.**
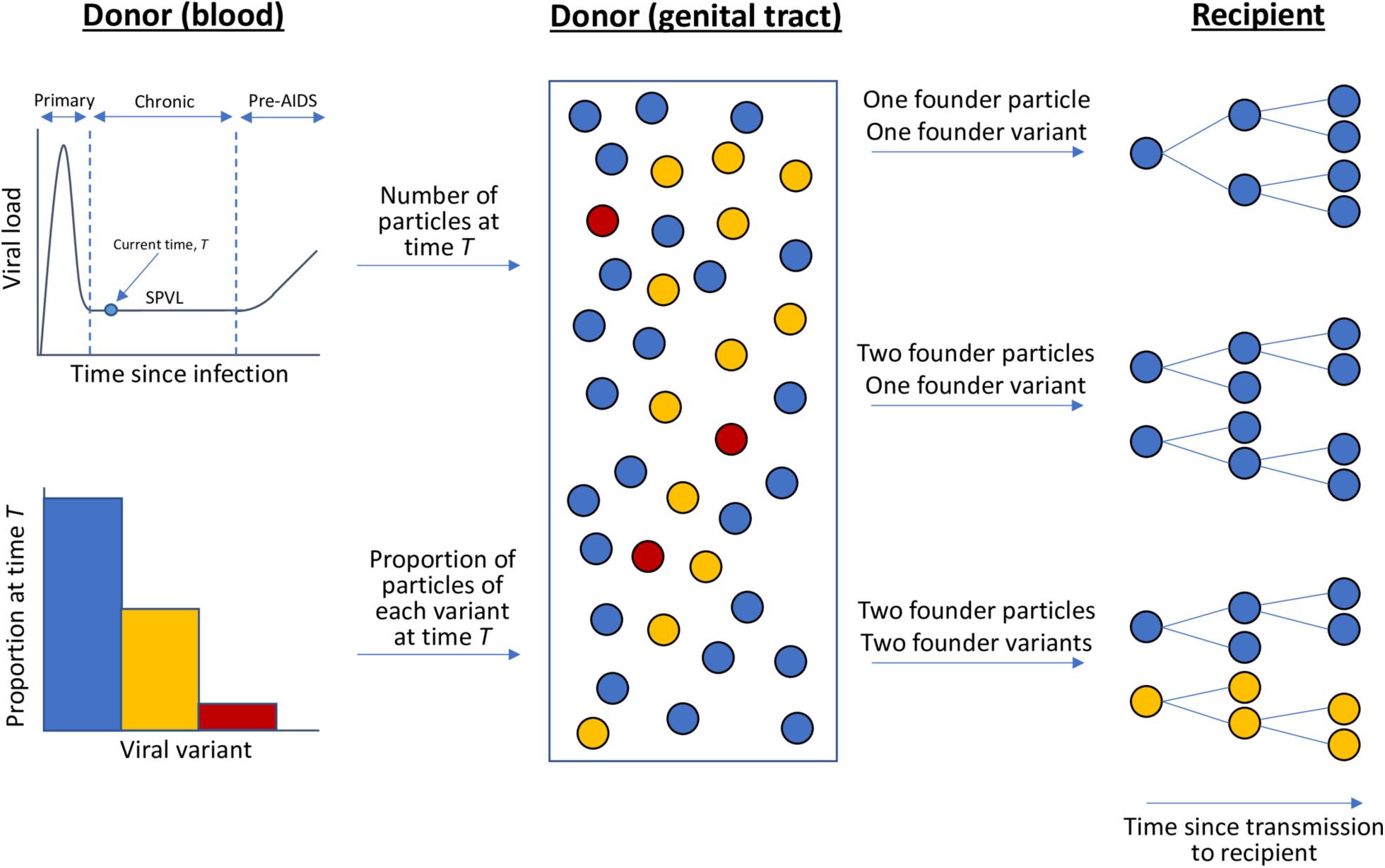
Schematic illustrating how the number of viral particles and number of variants founding a new infection were described by our model. The viral load and the proportion of each variant in the donor (potentially accounting for selection at transmission, as described in Materials and Methods) were used to determine the number of particles of each variant available for transmission in the donor’s genital tract. When a successful transmission act occurred, the particles that successfully founded the new infection were sampled at random from those in the genital tract.

To connect from this single transmission event scale to the population scale, we then considered a population of donors with different SPVLs (Figure 2A) and at different stages of infection. The composition of SPVLs in the donor population was determined using data on the proportions of infected individuals with different SPVLs within a population and a characterisation of their expected profiles of infection [21], both shown in Figure 2. The proportion of individuals with each SPVL is slightly different to the distribution described by Fraser *et al*. [21]: ours has been adjusted from data restricted to seroconverters to represent a full population of donors at different stages of infection. This reflects the fact that seroconverters who go on to have high SPVLs will survive for shorter periods than those with low SPVLs. Finally, we used previously published longitudinal deep-sequencing data to parameterise a function describing the expected distribution of unique variants within an individual throughout infection, as described below.

**Figure 2.**
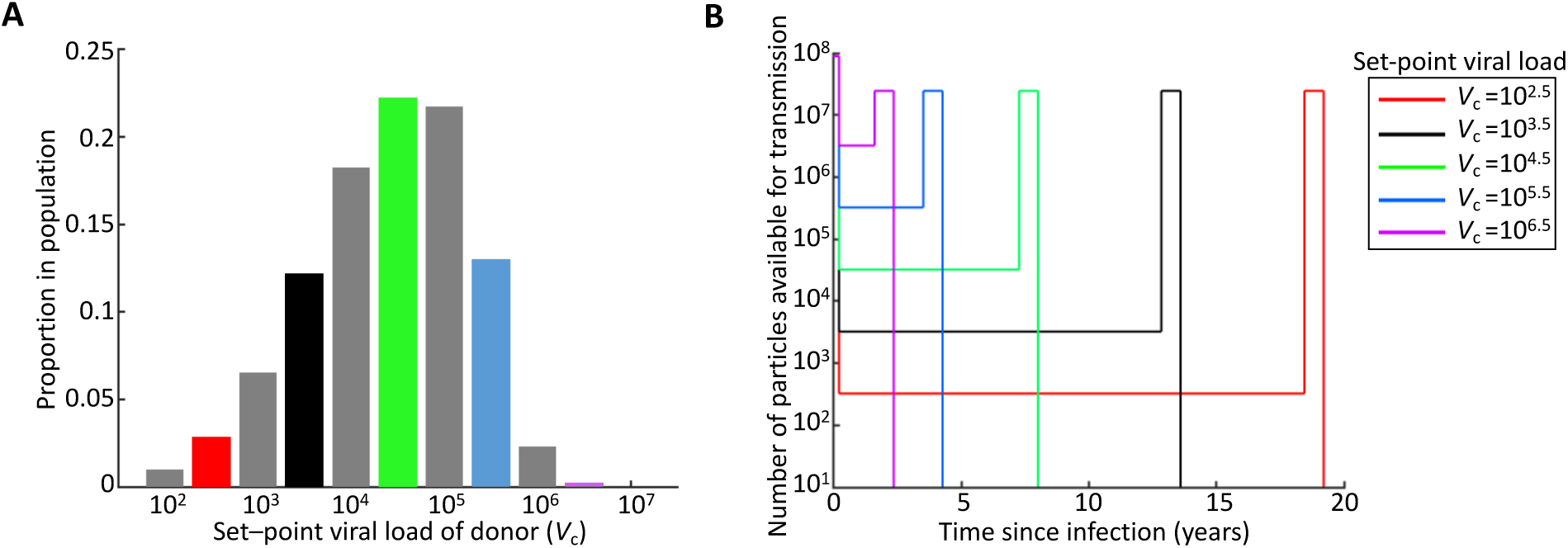
Viral loads in donors. (A) Proportions of infected individuals with different SPVLs. (B) Viral load profiles assumed in our model. Viral load profiles shown in B appear in the population of donors according to the proportions shown in A. Some of the SPVL values from A are omitted in B for clarity, but all values in A are included in our analyses.

### The distribution of viral variants as infection progresses

We fitted a nonlinear mixed-effects model to previously published whole-genome deep-sequencing data from longitudinally sampled infected hosts [18] to characterise the distribution of variants in an untreated individual as an HIV-1 infection progresses. In the absence of selection at the point of transmission, we assume this reflects the distribution of variants available for transmission in each individual in the population. For all three regions of the genome analysed (integrase, p24 and nef), a discretised gamma distribution provided the best fit to the data, characterising *h*(*x*,τ) – the proportion of the *x*^th^ most common viral variant in the within-donor pathogen population at time τ years since the individual became infected (see Materials and Methods). Since the highest viral diversity was observed for integrase, we used the parameterisation for this region in our main analysis (see Table 1 of Text S1). From the data, and our model fit, it can be seen that in the early years of an infection a small number of variants dominate, but as an infection progresses a higher diversity of variants (i.e. a more uniform distribution of variants) is seen (left column of Figure 3). Throughout our manuscript, by high diversity of variants we mean an approximately uniform distribution of variants as opposed to a distribution skewed so that there are high proportions of some variants and low proportions of others. The corresponding distributions for p24 and nef are show in Figure S1.

**Figure 3.**
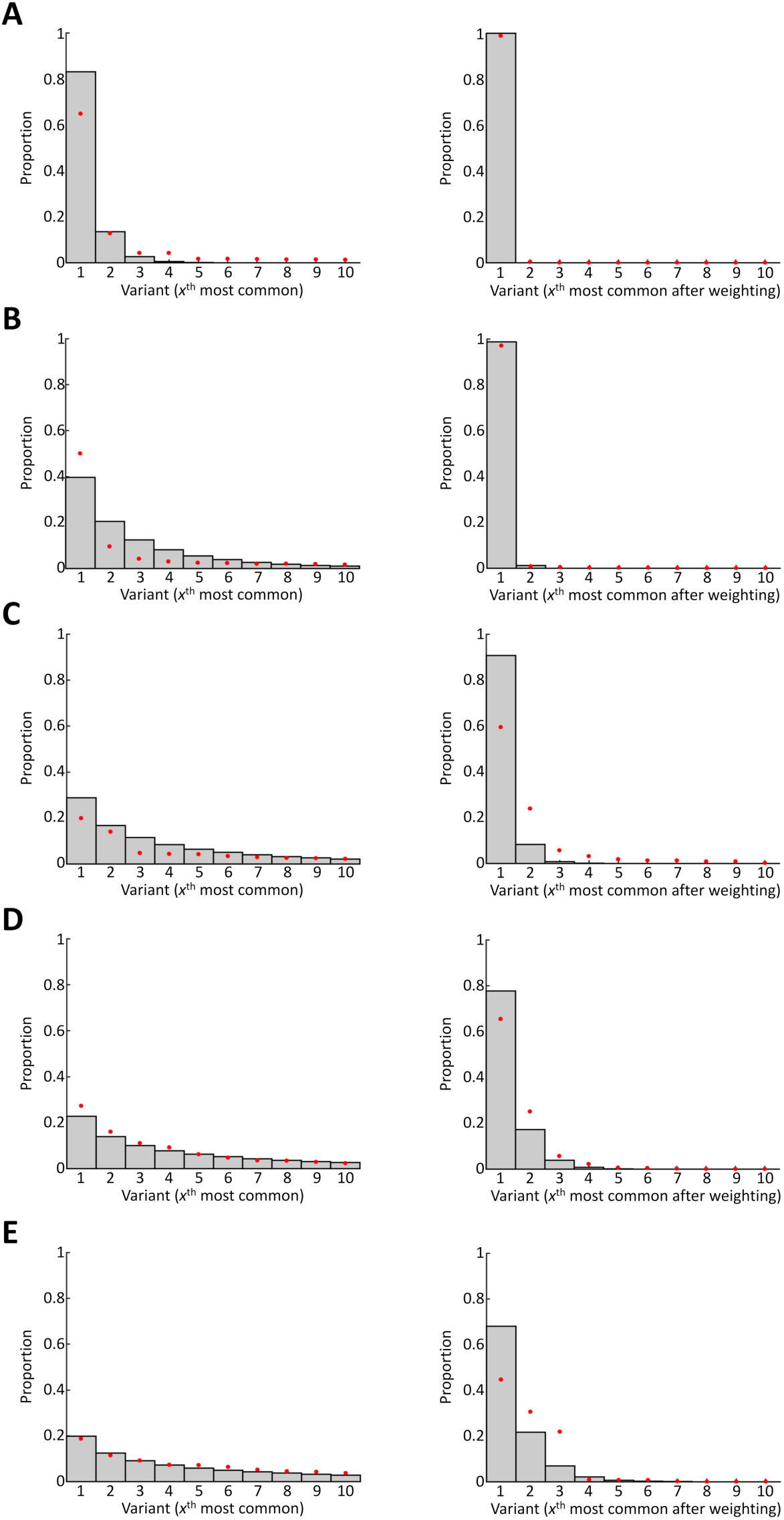
Distribution of variants in donors during the course of untreated infection. All data are for integrase. In the left column, the *x*-axis represents the *x*^th^ most common variant at the time of sampling, whereas in the right column the x-axis represents *x*^th^ most common variant after adjusting for selection (*α*_s_ = 3). Note that the *x*^th^ variant in one subfigure does not necessarily correspond to the *x*^th^ variant in another subfigure. Bars represent the model fit to all the data using a discretised gamma distribution (see Materials and Methods), and the red dots represent the data from one of the individuals (individual 3). We displayed data from this individual because of their long duration of infection and approximately equally spaced sampling time points. (A) 0.4 years since infection; (B) 2.2 years since infection; (C) 4 years since infection; (D) 6.4 years since infection; (E) 8.4 years since infection. Data from other infected individuals, and other regions of the viral genome, are shown in Figures S10-S12.

To incorporate selection at transmission into our analysis, we assumed that variants that are more similar to those that initiated the infection are more likely to be transmitted, since these represent variants that previously were successfully transmitted [28–30]. This assumption is supported by the faster rates of evolution of HIV-1 within-than between-hosts [28,31,31], the transmission of slowly evolving within-host lineages in a large transmission chain [33], and evidence for the transmission of founder-like virus in transmission couples [34,35]. We weighted the relative proportions of each variant in the sequencing data based on how close they are to the consensus sequence at the first time point in that donor, and then refitted our model (see Materials and Methods, and also Table 1 of Text S1). When selection is included, the effective diversity of variants available for transmission is reduced (right column of Figure 3). We show the very strong selection case here (*α*_s_ = 3), but results are also shown for strong and intermediate selection in Figure S2, where the parameter *α*_s_ is a measure of the strength of selection. The most common variant available for transmission in the presence of selection is not necessarily the most common variant in the absence of selection. Here we assumed that selection acts through the preferential transmission of founder-like variants, leading to a reduction in the diversity of variants available for transmission. However, any form of positive selection, in which some genotypes are favoured over others, could be implemented in our modelling framework.

### Numbers of particles and viral variants that successfully found new infections

Using our transmission model, we characterised the relationship between the numbers of T/F particles and the numbers of T/F variants in newly infected individuals within a population. We set the proportion of the time that the environment within an uninfected individual is appropriate for transmission (*f*) and the per-particle transmission probability in each act when the environment is appropriate (*p*) so that transmission occurred in three out of every 1000 transmission acts [22], and multiple variants founded 30% of new infections [2,4,4], although we also later show that our qualitative results are robust to reasonable variation around these values. In general, specifying the per-act transmission probability and the probability of multiple T/F variants uniquely determines *f* and *p* for a given distribution of variants within the donor population (Figure S3). The values of *f* and *p* used for each of the cases we considered are given in Table 2 of Text S1. The distributions of the numbers of T/F particles and variants in the recipient population were then derived analytically for three scenarios: no selection, selection at transmission, and transmission biased towards early infection but no selection.

#### No selection (Case 1)

The probability that a new infection is founded by *n* particles decreases as *n* increases, with a chance of approximately 40% that a single particle is transmitted, and 25% that two particles are transmitted. Similarly, the probability that *N* variants are transmitted also declines as *N* increases (Figure 4A, top left). It is not always the case that a large number of T/F viral particles and a large number of T/F variants coincide (Figure 4A, top middle). When the donor is in early infection (infected for less than two years), transmissions are more likely to be with multiple particles but few variants (Figure 4A top right) than in later infection. This result is driven by donors in the primary phase of infection, in which viral loads are high but viral diversity is low, leading to a high probability of transmission of multiple particles to a recipient, but with transmissions made up mostly of single variants (Figure 4A, bottom left). In chronic infection, viral loads are lower, and so almost all successful infections from a chronically infected donor are with a single viral particle and a single viral variant (Figure 4A, bottom middle). When donors are in the pre-AIDS phase of infection, new infections are often founded with an equal number of viral particles and variants (Figure 4A, bottom right). Infections founded with two particles are likely to be associated with two variants (Figure 4A, top middle). However, the importance of the stage of infection of the donor results in the counterintuitive observation that infections founded by three particles are more likely to be associated with only a single variant than with any other individual number of variants (Figure 4A, top middle). This is because three-particle infections are most likely to arise when the donor is in primary infection (Figure 4A, bottom left), so that few variants are transmitted, whereas infections with two particles often arise from donors in later infection when diversity is high (Figure 4A, bottom right). See Figure S4 for the figures equivalent to the top row of Figure 4 using the variant distributions parameterised for p24 and nef.

**Figure 4.**
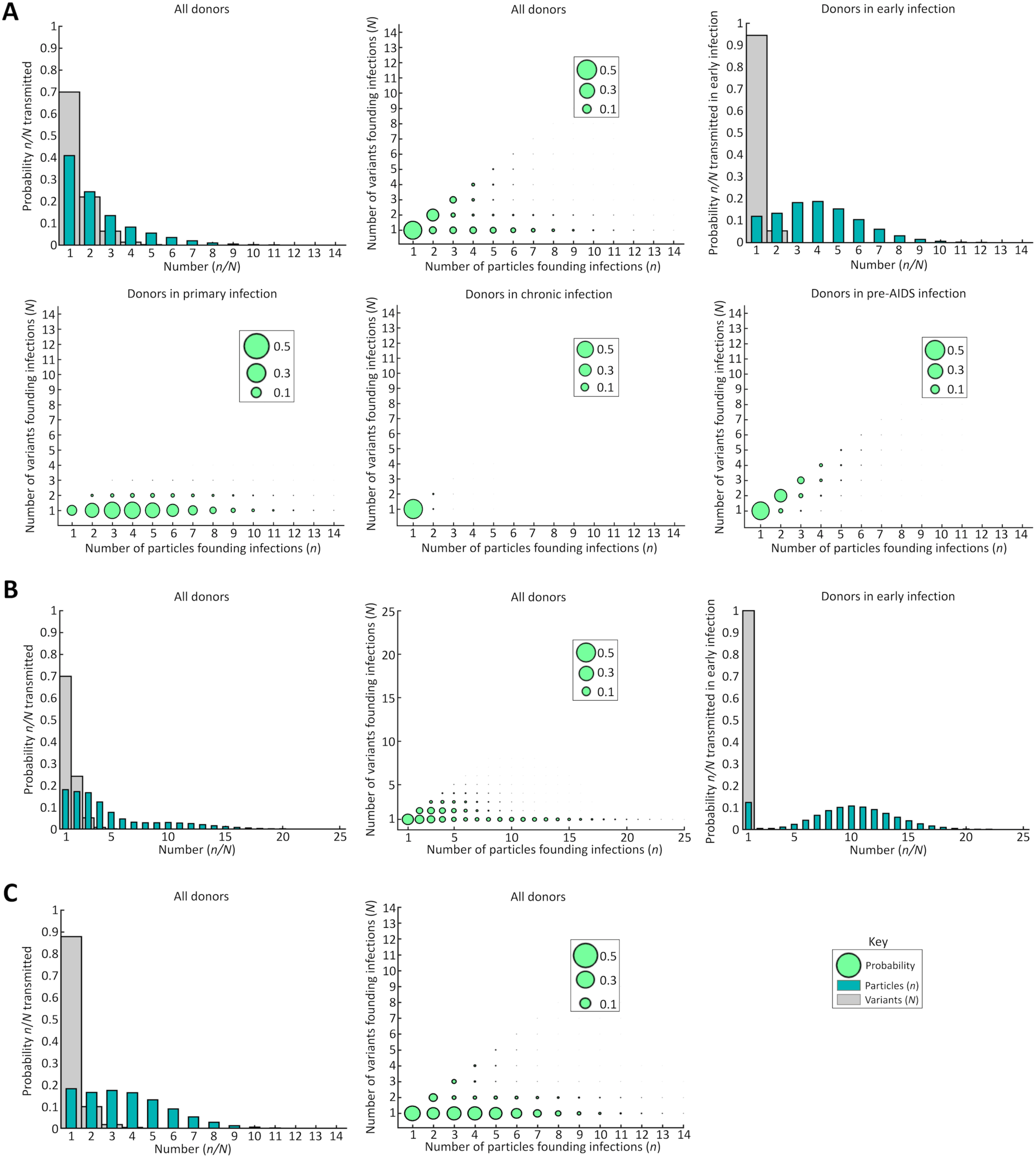
Distributions of transmitted numbers of particles and variants. (A) No selection and no weighting to early infection. The top left panel gives the distributions of the numbers of particles (teal) and numbers of distinct variants (grey) founding new infections in the population. The top middle panel shows the joint probability distribution of the numbers of particles and variants founding new infections; the area of each circle is proportional to the probability that exactly *n* particles and *N* variants were transmitted. The top right panel gives the distribution of the numbers of particles (teal) and numbers of distinct variants (grey) founding new infections in the population, from donors in early infection only (infected for less than two years). The bottom panels are the joint probability distributions that *n* particles and *N* variants are transmitted, conditioned on the donor being in primary (bottom left), chronic (bottom middle) and pre-AIDS (bottom right) infection. (B) Figures analogous to the top row of A, with selection at transmission (*α*_s_ = 3). (C) Figures analogous to the top row of A, with no selection at transmission but with a bias towards early transmission. The right panel is omitted since the bias towards early infection is assumed to change the proportion of infections in early infection (before 2 years), but not the composition of transmissions occurring during early infection, and so the result is identical to the top right panel of A. In the case with selection, due to the reduced diversity of variants available for transmission, some infections must be with large numbers of particles so that Prob(transmit multiple variants) = 0.3. Because of these large numbers of particles, the ranges on the x-axes in all panels of B and the y-axes of the middle panel of B are larger than in the equivalent subfigures in A and C. For parameter values, see Tables 1 and 2 of Text S1. In C, the same parameter values as A were used but with infection *w* = 10 times more likely at times when donors have been infected for less than τ_crit_ = 2 years.

By fitting a single distribution to the sequencing data from Zanini *et al.* [18], we effectively made the simplifying assumption that every donor had an infection that was originally founded by the same number of variants. Most of the individuals in that cohort were probably infected by single variants [19], and the fitted distribution of variants reflected this. Since around 30% of donors in real populations would instead have been infected by multiple variants, we conducted a supplementary analysis in which we assumed that 30% of donors had infections founded by two distinct variants, and that the resulting lineages from each T/F variant evolved independently within an individual (Text S1 – Multiple variants founding infections in donors). The results were qualitatively similar: the link between the numbers of T/F particles and T/F variants depended on the timing of transmission. Since we fitted a single distribution to the sequencing data, we also assumed implicitly that the distribution of variants in a donor is independent of SPVL. This is supported by longitudinal sequencing data in which a link between SPVL and viral diversity is not apparent [19,36].

#### Selection at transmission (Case 2)

We investigated how selection for particular variants would affect our results, and how sensitive this is to the strength of selection, *α*_s_. We used the fitted distributions of variants available for transmission with selection, as described above (see right panel of Figure 3 and Figure S2). Using these new distributions of variants, we reparameterised the values of the per-particle transmission probability (*p*) and the proportion of the time the environment is appropriate for transmission (*f*) so that the probability of transmission occurring per act remained at 0.003 and the probability of multiple variants founding each new infection was 0.3 (see Table 2 of Text S1). We carried out this reparameterisation step because we sought to consider the numbers of transmitted particles and variants if selection is currently acting in heterosexual populations for which transmission occurs in three out of every 1000 potential transmission acts and 30% of infections are founded by multiple variants. If selection had instead been imposed without reparameterising the model, then the numbers of transmitted variants would have been reduced compared to the case with no selection.

Even for very strong selection (*α*_s_ = 3), since we reparameterised *f* and *p* we found that the overall distribution of the numbers of variants founding new infections remained similar to the case in which there is no selection at transmission (Figure 4B; see Figure S5 for equivalent figures with different strengths of selection). However, because selection reduces the diversity of viral variants available for transmission (right column of Figure 3), a higher per-particle probability of transmission per act (*p*) was required to achieve 30% of new infections being founded by multiple variants. As a consequence, it became more likely that many particles were transmitted compared to the case in which there is no selection, but still only one or a few variants (Figure 4B). In other words, in terms of the numbers of transmitted variants, the reduced diversity of variants available for selection was cancelled out by the larger numbers of particles likely to be transmitted (which permitted more variants to be transmitted). Since large numbers of particles yet few distinct variants could then be transmitted, including selection in the model enhanced the prediction that the numbers of particles and variants founding new infections are not closely linked quantities.

#### Bias towards early transmission (Case 3)

There are many reasons why there might be a bias towards early transmission. Interventions are likely to lead to a bias towards early transmission, because awareness of HIV-1 status can cause behaviour changes and treatment reduces infectiousness [37–39], but there is a delay between infection and diagnosis and a further delay before action is taken. High rates of transmission due to needle sharing when injecting drugs [40,41] or concurrent partnerships [42,43] might also increase the number of contacts an individual has during the highly infectious primary stage of infection [44].

We therefore considered the distributions of transmitted particles and variants when transmission is more likely in early infection than in later infection. We assumed that potential transmission acts occur in early infection (defined to be time since infection τ < τ_crit_ years) at a rate enhanced by a factor *w* > 1 compared to later time points (see Materials and Methods). Since our model was parameterised using population-level data in which it is assumed that there is no bias towards early transmission, we did not change the values of *f* and *p* here from the no selection case considered above.

In Figure 4C, we considered the case where *w* = 10 and τ_crit_ = 2 years, representing for example a population in which test and treat interventions are very effective. A greater proportion of new infections were derived from donors in early infection than in the absence of bias towards early transmission, and so transmissions consisted of fewer distinct viral variants, but larger numbers of particles per successful transmission act. When a smaller value of the weighting parameter, *w*, was used, a similar but less extreme pattern was seen (Figure S6).

### Link between donor SPVL and recipient number of founder variants

We also considered the characteristics of the donors in the population that were most likely to transmit multiple variants (Figure 5).

**Figure 5.**
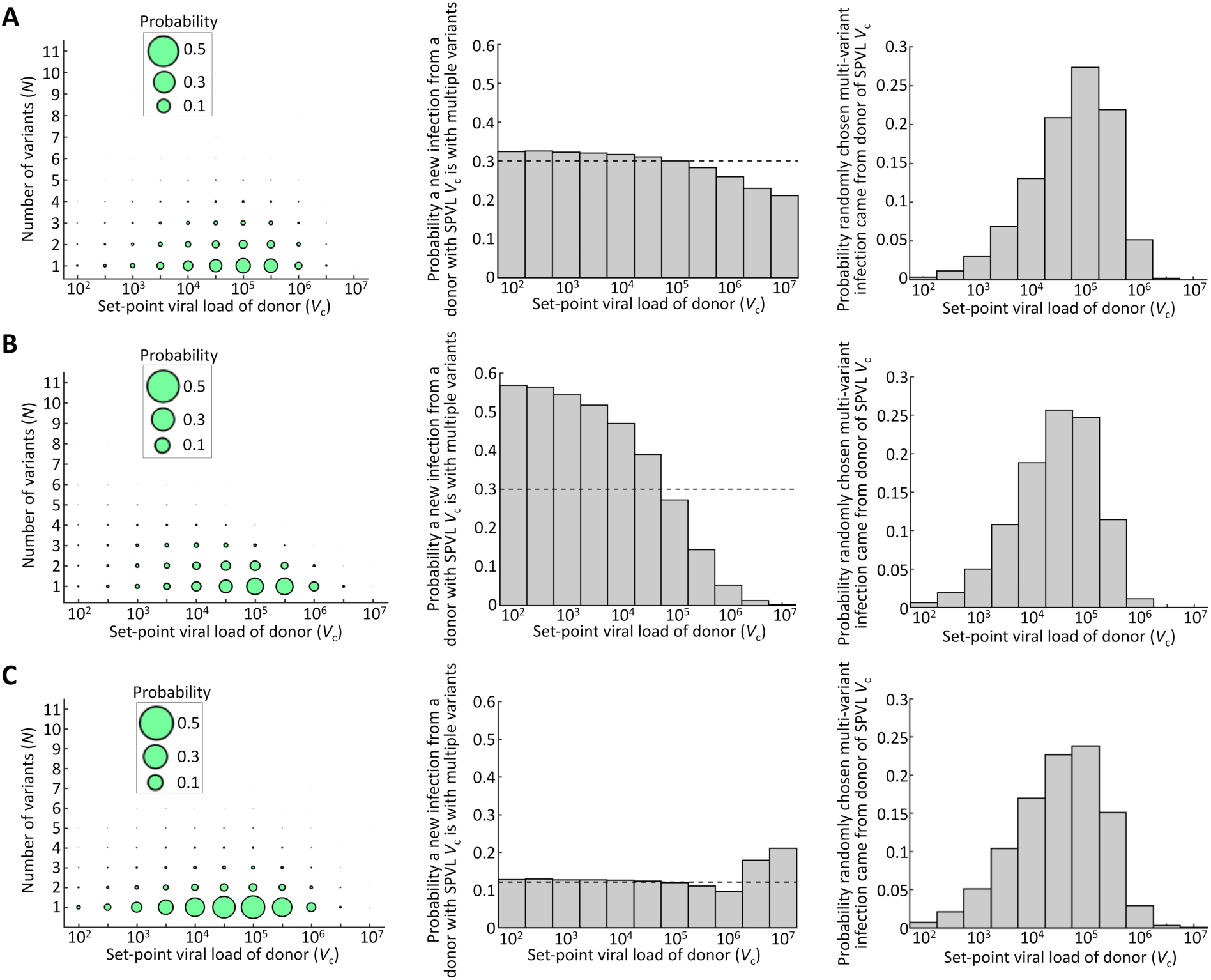
Relationship between number of T/F variants and donor SPVL. (A) No selection and no weighting to early infection. Left: The joint probability distribution of the number of distinct variants initiating a randomly chosen infection and the SPVL of the donor. The area of each circle is proportional to the probability of randomly chosen infection both originating from a donor with SPVL *V*_c_ and being founded by *N* variants. Centre: Conditional on transmission, the probability that an individual with SPVL *V*_c_ transmits multiple variants. Right: Probability that a randomly chosen infection initiated by multiple variants arose from a donor with SPVL *V*_c_. (B) Figures analogous to A, but with selection at transmission (*α*_s_ = 3). (C) Figures analogous to A, but with a bias towards early transmission. For parameter values, see Tables 1 and 2 of Text S1. In C, the same parameter values as A were used but with infection *w* = 10 times more likely at times when each donor has been infected for less than two years.

#### No selection (Case 1)

We derived the joint distribution characterising the numbers of transmitted variants and the SPVLs of donors in the population (left panel of Figure 5A). The most likely combination was a single variant infection arising from a donor with intermediate SPVL, reflecting the fact that most infections are with single variants (Figure 4A), and most infected individuals have intermediate SPVLs (Figure 2A).

The chance that a randomly chosen infection from a donor with each SPVL consisted of multiple variants is shown in the middle panel of Figure 5A. It can be seen that, despite donors with high SPVLs being likely to transmit large numbers of particles, each infection is likely to be with few variants. This is because donors with high SPVLs are likely to die more quickly than individuals with lower SPVLs, so they tend to transmit before the founder viruses have diversified.

We then focussed solely on infections arising with multiple T/F variants. High-SPVL donors are not only uncommon, but also tend to transit only one variant due to their short duration of infection, meaning only a small proportion of infections founded by multiple variants arose from donors with high SPVL. Compared to donors with intermediate SPVL, low-SPVL donors are also uncommon, and not very infectious, which together outweigh the fact that their long duration of infection provides more time for the virus to diversify. Consequently, most infections in the population that were founded by multiple variants arose from donors with intermediate SPVLs (right panel in Figure 5A). Some of the intermediate quantities for calculating the results of Figure 5A are shown in Figure S7.

#### Selection at transmission (Case 2)

When selection is incorporated into the model, a donor with a low SPVL is now much more likely to produce a multiple variant infection than a donor with higher SPVL (centre panel in Figure 5B). This is because including selection makes it more likely that transmission early in infection will result in new infections founded by only one variant. As a result, the donors who survive for long periods have the opportunity to transmit after viral diversity has increased, and these are the donors with low SPVLs. In the selection case, we therefore find that infections founded by multiple variants are likely to have come from donors with lower SPVLs than the case with no selection (Figure 4B right panel).

#### Bias towards early transmission (Case 3)

When transmission is heavily weighted towards early infection, individuals with very high SPVLs become more likely to transmit multiple variants than individuals with low SPVLs (Figure 5C middle). When there is no bias towards early transmission, high-SPVL individuals have shorter durations of infection than other individuals, and so have less opportunity to transmit after viral diversity has accumulated. When there is a strong bias, however, all individuals effectively have similar, shorter, durations of infectiousness, and are all unlikely to transmit before the founder viruses have diversified significantly. In this case, the increased probability of transmitting multiple viral particles (and so multiple variants) at higher SPVLs becomes more important. However, even in this case, a randomly chosen multiple-variant infection is most likely to have originated from a donor with intermediate SPVL (Figure 5C right), since most donors have intermediate SPVLs.

## DISCUSSION

We developed a probabilistic model to characterise HIV-1 transmission in an untreated population, focussing on the relationship between the numbers of particles and the numbers of variants founding new infections. A key finding from the model is that transmissions with more T/F particles are not necessarily those with more T/F variants, with the timing of transmission during the donor’s course of infection being of critical importance. This is especially noticeable when donors transmit during primary infection, since viral loads are high but viral diversity is low. The observation that most infections are initiated by one or a few variants has been used as evidence for the role of selection at the point of transmission and/or during establishment of infection in recipients [1,9,9]. In particular, if a new infection is founded by few variants, then it might be assumed that selective factors, such as physical barriers to transmission, are preventing other variants from being transmitted or successfully establishing the infection in the recipient. However, the role of selection, as opposed to transmission simply being a stochastic process, has been debated [11–15].

Here, we have shown that by considering viral diversity within donors explicitly, imposing selection is not required to reconcile within-host and population-level data, or to explain the low numbers of T/F variants generally observed. We do not contend that selection is unimportant - a number of phenotypic transmission factors have been identified (e.g. [1,44–46]) - but rather that the viral bottleneck at transmission is likely to be due to both selective and stochastic forces. By including selection in the model, we found an even weaker link between the numbers of T/F variants and T/F particles than we found in the absence of selection, with a higher proportion of infections being founded by large numbers of particles but few variants. Similarly, when a bias towards early transmission was included in the model, which could be due to treatment or other behavioural changes, single-variant but multiple-particle transmission became more likely.

We have shown that the distribution of viral variants during the course of infection, the timings of transmissions, and the strength of selection are some of the many factors that are likely to influence the compositions of new HIV-1 infections. This complex interaction of different factors could help to explain some confusing observations. For example, Tully *et al.* [8] found that the proportion of infections founded by single variants in an MSM population was similar to that typically observed in heterosexual populations, despite finding signatures of reduced selection compared to heterosexual transmission. This is puzzling because reduced selection is expected to result in more variants being transmitted, since more variants are likely to possess characteristics that permit successful transmission and establishment of a new infection. This can be explained if transmission tended to occur early in the MSM population considered by Tully *et al.* [8], as has been suggested for MSM transmission more generally [42,43]. This is because lower viral diversity within the pool of transmitting donors due to early transmission will tend towards more new infections being founded by single variants. The increased diversity of infections due to reduced selection may therefore have been balanced by the reduced diversity of infections due to early transmission. Given these complex interactions, we urge caution in interpreting the proportion of infections founded by single variants as a universal statistic, even between populations that on the surface appear quite similar. Differences in the timing of transmission between different MSM populations might help to explain the higher proportion of infections founded by multiple variants observed in some studies [2,47,47] compared to others [3,6–8], although differences in sequencing and methods of analysis might also have a role [48]. All else being equal, we would predict fewer variants being transmitted in populations where transmission is biased towards early infection.

A positive association between multiple variants founding an infection and a high SPVL of that infection has been observed [16]. Since a higher SPVL is associated with faster progression to AIDS, understanding the factors leading to multi-variant transmission is important for inferring the mechanisms driving the severity of different HIV-1 infections. This will also inform the development of evolutionary epidemiological models. It has been suggested that recipient host factors might be important [1,16]. We hypothesised that the SPVL of the donor might be another key factor involved in multi-variant transmission. Contrary to what might be expected, we found that most infections founded by multiple variants do not arise from donors with high SPVLs, but from donors with intermediate SPVLs; individuals with high SPVLs tend to rapidly progress to AIDS, and therefore viral diversity has limited time to accumulate. It is not known why there is a positive association between multi-variant transmission and a higher SPVL, although it has been suggested that viral diversity *per se* has a role [16,48]. Another possibility is that when more variants are transmitted, there is a higher chance that one of these variants possesses viral factors associated with high SPVLs within the recipient.

Our aim here has not been to develop a detailed model of HIV-1 transmission, but rather to present the simplest possible model that encapsulates important features of transmission within a population. We therefore made a number of simplifying assumptions, including assuming random contacts between donors and potential recipients, and ignoring host genetic factors that might affect viral diversity within donors.

Nonetheless, most other simple models cannot accommodate the infrequent transmission, yet reasonably high proportion of infections founded by multiple variants, observed in real populations [4]. We captured this by assuming that transmission is only possible a small fraction of the time, when the environment is appropriate. In doing this, we assumed that when a potential transmission act occurs the environment is either entirely permissive (each available viral particle can be transmitted independently of the others with a constant probability) or entirely resistant to transmission. In reality, the permissiveness of the environment to transmission is likely to be a continuous quantity, rather than always entirely “on” or “off”. Abrasions in the genital tract, genital inflammation or infection with other pathogens might increase the probability of transmission [12,23–27]. Facilitation, whereby a virus being transmitted changes the environment in the recipient so that further transmission is more likely to immediately occur [4,49], might also be able to reconcile infrequent transmission with reasonably frequent multi-variant transmission. In the facilitation scenario, the probability of transmission is assumed to be low, yet when a particle is transmitted the environment temporarily changes enabling further particles, and therefore potentially multiple variants, to be transmitted. Other mechanisms might also be able to reconcile the low transmission probability of HIV-1 with the significant proportion of new infections founded by multiple variants, and could provide an interesting avenue for further exploration using theoretical models.

The link between viral load and the transmission rate is also in need of further study [50]. Results from models based on binomial distributions for the numbers of transmitted particles, such as the model we have developed, display an approximately linear relationship between the viral load and the transmission rate. In contrast, empirical evidence suggests that the transmission rate increases linearly with the logarithm of the viral load [21,51]. This discrepancy might arise partly because the relationship between the SPVL and the transmission rate has been determined in monogamous heterosexual couples, making it difficult to detect multiple transmissions from the same donor and thus underestimating rates of transmission when viral loads are high. Assuming a nonlinear relationship between the viral load and the number of particles available for transmission in the donor’s genital tract and/or a nonlinear relationship between viral load and viral fitness [52] might also allow binomial models to reproduce observed data.

A key measure that we have approximated in our model is the distribution of viral variants within donors as infections progress. We used previously published short-read deep sequencing data from longitudinally sampled untreated individuals [18] to estimate the distribution of distinct variants in a typical donor throughout infection. However, the diversity of variants estimated using a short segment of the genome is almost certainly going to underestimate the true diversity [53–56], and conversely, sequencing errors are likely to lead to an overestimate of the number of rare variants [57]. As a result, our fitted distributions characterising the diversity of variants in donors might not reflect the diversity present in a typical individual throughout infection. To investigate the importance of this uncertainty for our model predictions, we repeated our analyses assuming both lower and higher variant diversities within donors by varying the parameters of the gamma distribution characterising variant diversity (Figure S8), as well as different values of the per-particle transmission probability (Figure S9). By varying the per-particle transmission probability in Figure S9, we also implicitly tested the robustness of our results to the assumption that 30% of new infections are founded by multiple variants. Our key conclusion remained unchanged: the link between the numbers of T/F particles and variants depends on the timing of transmission.

Understanding the relative roles of selection and other factors in determining the strong bottleneck that occurs during HIV-1 transmission is relevant for vaccine design [58] and determining the drivers of pathogenesis [59], as well as in the development of epidemiological and phylodynamic models that can capture viral transmission in a realistic fashion. Here, we have highlighted the need to consider viral diversity in donors at the times of transmissions as an additional important, but hitherto underappreciated, factor.

## MATERIALS AND METHODS

### Modelling temporal changes in donor viral load

Following a previously used modelling approach [21], we divided the infectious period of an infected donor into three stages: primary, chronic and pre-AIDS. The viral load of the donor depends on the stage of infection. For the full mathematical details of this approach and its parameterisation, see Text S1 – Viral load profiles; a summary is below.

In primary infection, which lasts τ_p_ = 0.24 years, the viral load for all donors is *V*_p_ = 8.7 x 10^7^ viral particles per millilitre of blood. During the chronic stage of infection, the viral load *V*_c_ is fixed at set point, which varies by several orders of magnitude between donors [60,61]. Donors with higher SPVLs progress to AIDS more quickly than individuals with lower SPVLs [21], so that the time spent in chronic infection τ_c_(*V*_c_) depends on the SPVL. The probability that a randomly chosen donor has SPVL equal to *V*_c_, which we denote g(*V*_c_), is shown in Figure 2B and given in detail in Text S1 – Viral load profiles. In the pre-AIDS stage of infection, which lasts τ_a_ = 0.75 years, the viral load is *V*_a_ = 2.4 x 10^7^ viral particles per millilitre for all individuals.

Throughout infection, the number of particles available for transmission in each donor in the population is assumed to be proportional to the viral load. We denote the number of particles available for transmission in donors by *n*_p_, *n*_c_ and *n*_a_ in primary, chronic and pre-AIDS infection, respectively. In the analyses in the main text, we have assumed that the constant of proportionality, *k*, is equal to one, so that e.g. *n*_c_ = *V*_c_. We note that our results are very similar if different values of *k* are used. This is because the results depend (approximately) only on the product of *k* and the per-particle transmission probability, *p*, rather than the individual values of these parameters – and larger values of *k* correspond to smaller values of *p* when the model is refitted so that the per-act transmission probability is 0.003 (see Text S1 – Relationship between viral load and number of particles available for transmission).

### Modelling variant diversity in donors

We used publicly available whole-genome deep sequencing data from ten longitudinally sampled HIV-1 infected individuals (individuals 1-3 and 5-11 from Zanini *et al*. [18]) to obtain an approximation of the distribution of HIV-1 variants within a typical infected individual as infection progresses.

Specifically, we used information from three distinct regions of the viral genome, chosen for their wide coverage and because they come from three different functional categories: integrase (enzyme, HXB2 reference positions 4230-5096); p24 (structural, positions 1186-1878); and nef (accessory, positions 8797-9417). Each of the reads from these regions are around 300 base pairs long and we conducted our analyses separately for each region. We only included samples that contained a large number (at least 1000) of reads, so that a distribution of variants within each sample could be characterised.

We assumed that each distinct read corresponded to a different variant of the virus, and then found the proportion of each variant in each sample. Variants at proportions lower than 0.005 were removed, to protect against sequencing error. The resulting distribution of variants in one of the individuals (individual 3) throughout their course of infection, obtained using data from integrase, is shown by the red dots in Figure 3. The distributions of variants obtained from the remaining individuals, and for the different regions of the viral genome, are shown in Figures S10-S12.

To characterise *h*(*x*,τ) – the proportion of the *x*^th^ most common variant in each donor at time τ years after they became infected – we considered three candidate distributions: gamma, exponential and Pareto. These distributions all provided a reasonable fit to the data (see e.g. Figure 3). The probability distribution functions are

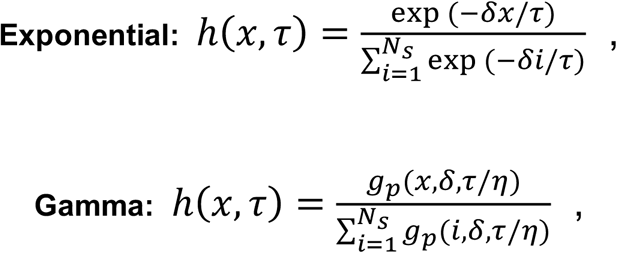

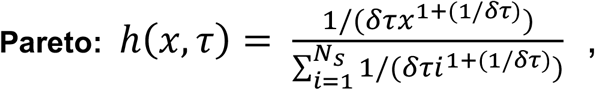

for *x* = 1,2,3,…*N*_s_, where *N*_s_ is the maximum number of distinct variants observed in any individual at any single time in the data. The values of *N*_s_ for integrase, p24 and nef were 56, 45 and 51 respectively. The function *g*_*p*_(*i,j,k*) is the probability that a gamma distributed variable with shape parameter *j* and scale parameter *k* takes the value *i*, i.e.

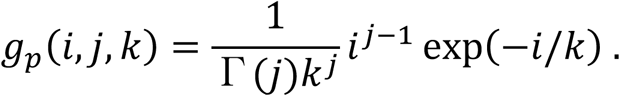

We fitted the parameters of each of the candidate functions *h*(*x*,τ) to data using a nonlinear mixed-effects model. This fitting was performed using the R software function *nlme* with fixed effects of variant *x* and time since infection τ, and a random effect of the individual that each read was sampled from. Including a random effect of the sampled individual amounts to a partial pooling of the data between individuals to improve our estimates of the parameters applicable to the broader population from which these individuals were drawn. By doing this, the differences between these individuals (which are not of direct interest, and are difficult to infer for individuals with little data) were not estimated, nor did we fully pool the data (which would bias estimation towards over-sampled individuals). The candidate models were compared using the Akaike information criterion scores associated with their model fits. The resulting parameter values for each of the three regions that we considered are shown in Table 1 of Text S1. While the gamma distribution that provided the best fit to the data has the property that there is effectively only a single variant available for transmission in the donor at small times since infection, we also considered cases in which there could be multiple variants at equal frequency early in a donor’s course of infection (Text S1 – Multiple variants founding infections in donors).

### Modelling selection

When modelling selection at transmission, we assumed that the variants most similar to those that donors were themselves infected with were more likely to be transmitted, since they are likely to retain characteristics that make them suited for onward transmission [30]. To investigate the effect of preferential transmission of these ‘founder-like’ variants on the distributions of the numbers of transmitted particles and variants according to our model, we manipulated the variant proportions in the sequencing data to obtain *effective* variant proportions, assuming a selection coefficient that decays exponentially with Hamming distance from the founding consensus sequence [29].

First, the founder sequence was estimated for each donor by taking their earliest available sample, and finding the most common base at each position. Then, if the proportion of sequence *x* in a donor in a reading taken at time *τ* years since infection is *h*(*x,τ*), where here we use *h*(*x,τ*) to refer to the proportion in the data rather than the fitted model, then the effective proportion in the donor was assumed to be

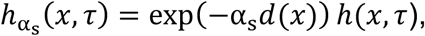

where the parameter *α*_s_ characterises the strength of preferential transmission of founder-like variants, and *d(x)* is the number of base pairs at which the variant *x* differs from the founder sequence for that donor. For example, for strength of selection *α*_s_ = 1, then the effective proportion of a variant with three base pairs difference from the founder is calculated by reducing the original proportion by factor exp(−3). The resulting distribution of variants in a particular donor at each time *τ* years since infection was then normalised so that the effective proportion of variants made up a valid probability distribution (for example if all variants are equally different from the founder, the effective distribution is identical to the original distribution without selection). Since the most common variant in the absence of selection was not necessarily the most common variant once selection had been applied, the variants were renumbered so that the variant with the highest effective proportion was labelled variant 1, and so on.

As in the case with no selection, the models were then fitted to the resulting distribution of the effective quantities of each variant. The best-fitting model and parameter values for the data from the integrase region are shown in Table 1 of Text S1 for *α*_s_ = 0,1,2,3. The value *α*_s_ = 0 corresponds to the case in which there is no selection.

### Modelling transmission

We assumed that environmental conditions are suitable for transmission in a fraction *f* of transmission acts, and that when conditions are suitable each particle has a probability *p* of being transmitted, independently of the other particles. The probability of *n* particles being transmitted and going on to generate a new infection is therefore given by

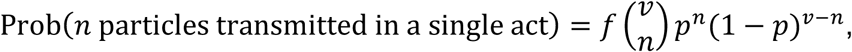

for *n* = 1,2,3…, where the parameter *v* is the number of particles available for transmission in the genital tract of the donor at the time that the potential transmission act occurs.

The probability of transmitting *N* distinct variants in any single potential transmission act is given by

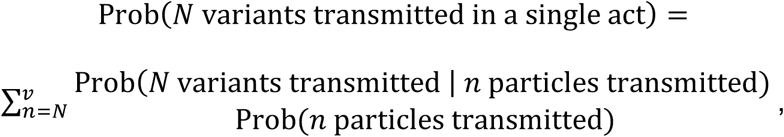

where the first factor in the sum depends on the effective distribution of variants available for transmission in the donor (accounting for changes in the effective diversity of variants likely to be transmitted given selection at transmission).

### Population-scale quantities

The following quantities were derived analytically by integrating over all infected potential donors in the population, and all times during their courses of infection. The variables *n*_*p*_, *n*_*c*_, and *n*_*a*_ represent the numbers of particles available for transmission in the genital tracts of donors in primary, chronic and pre-AIDS infection, respectively. In the version of the model described here, the number of particles available for transmission is equal to the viral load, so that for example *n*_*c*_ = *V*_*c*_ (*cf.* Text S1 – Relationship between viral load and number of particles available for transmission). The variables *τ*_*p*_, *τ*_*c*_(*n*_*c*_), and *τ*_*a*_ are the durations of infection in primary, chronic (for a donor with *n*_*c*_ = *V*_*c*_ particles available for transmission in chronic infection) and pre-AIDS infection, respectively. For n ≥ 1 and N ≥ 1:

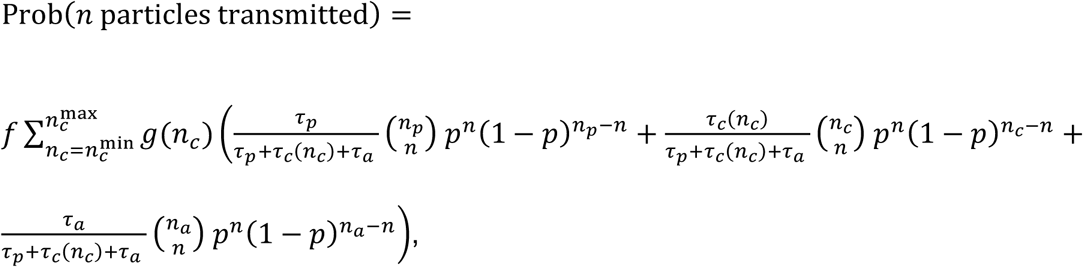

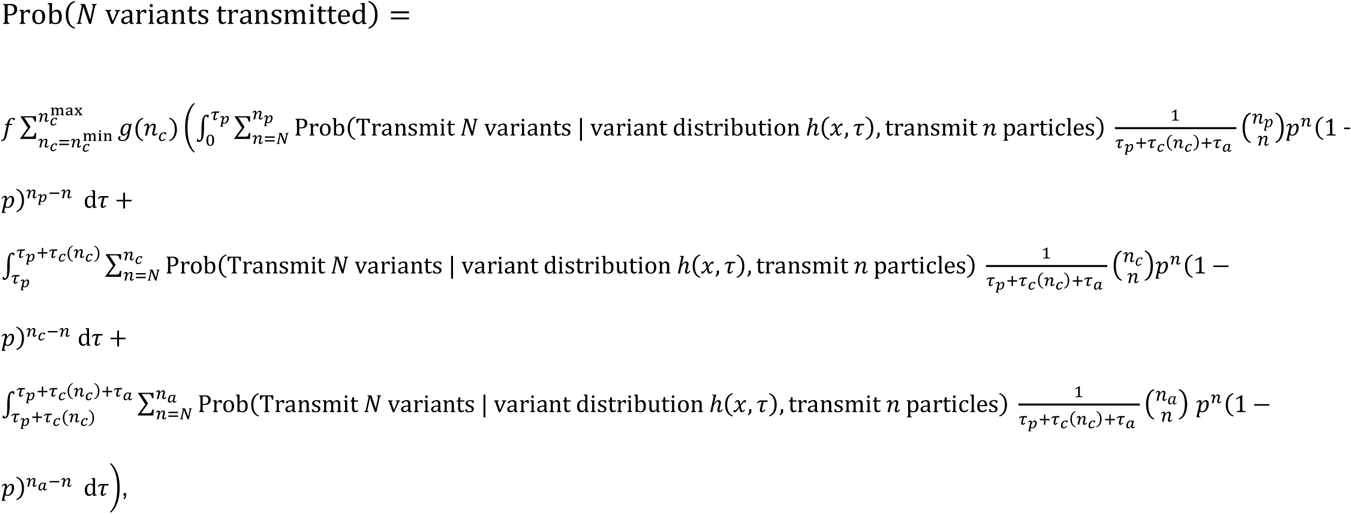

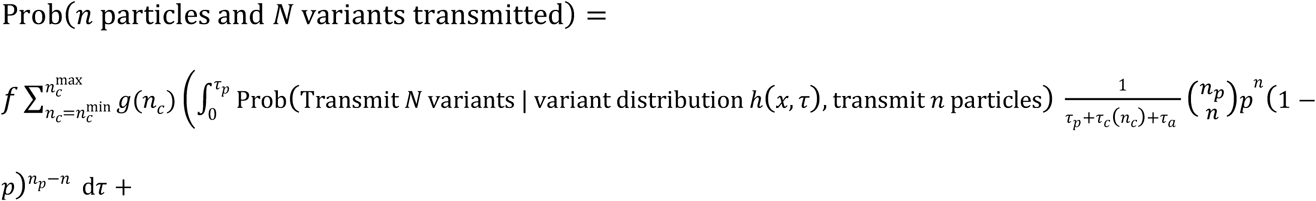

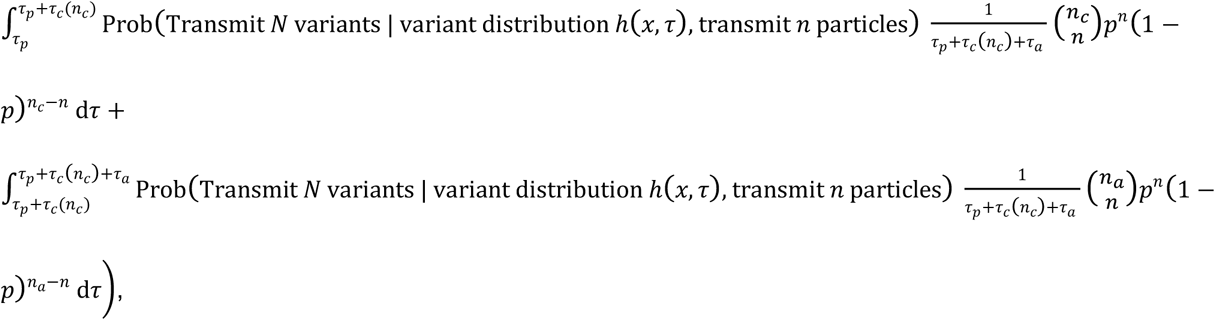

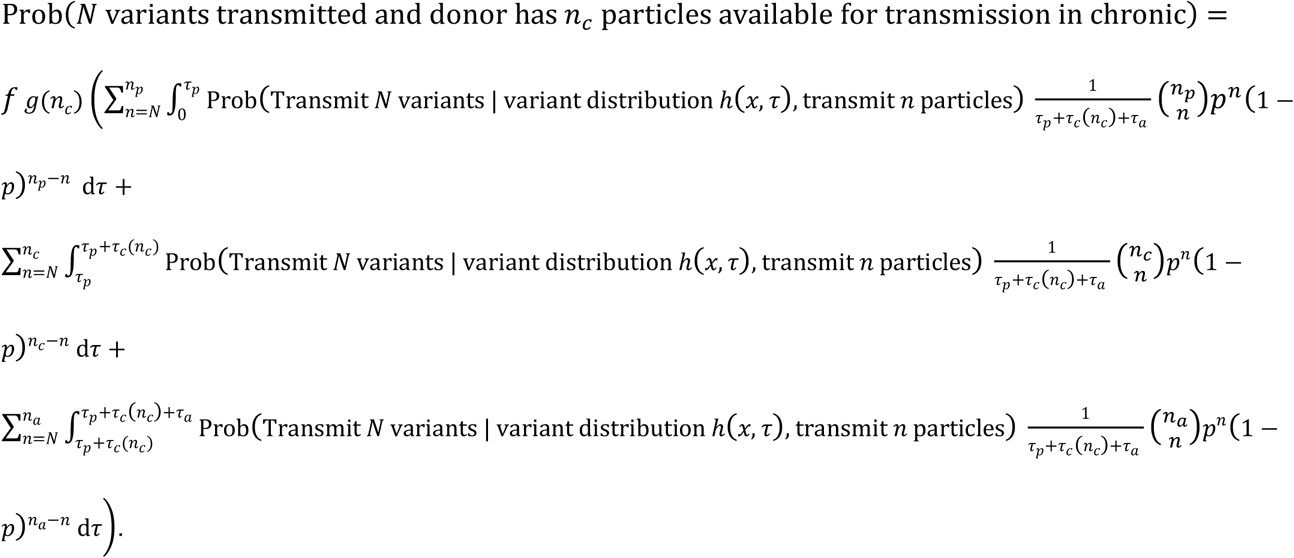

When we parameterised the model so that 30% of new infections are founded with multiple variants, we chose the per-particle transmission probability *p* so that 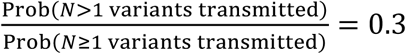. With this value of *p*, the value of the parameter *f* can be chosen so that three out of every 1000 potential transmission acts lead to successful transmission and establishment in the recipient by setting Prob(*n* ≥ 1 particles transmitted) = 0.003. This fitting procedure is shown in Figure S3. The expressions above were then normalised to obtain the probability distributions conditional on transmission occurring and successful establishment in the recipient.

### Modelling bias to early infection

If some individuals in the population are undergoing antiretroviral therapy, then transmission is more likely to occur when a donor is in early infection than in later infection once treatment may have started. As described in the Results, transmission might also be more likely in early infection for other reasons. We therefore considered populations in which the timings of transmissions are weighted towards early infection, in addition to the increased probability of early infections due to the higher viral load in the primary phase. As an example of how we implemented this, the 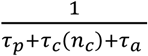 terms in the expression for Prob(*n* particles and *N* variants transmitted) were replaced by appropriate terms. If, for example, early infection is defined to be τ < τ_crit_ where τ_crit_ < τ_p_ + τ_c_(*n*_c_) + τ_a_, and the weighting factor is *w >* 1, then this term would be 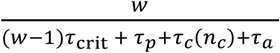 for τ < τ_crit_ and 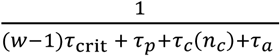 for τ > τ_crit_

## ACKNOWLEDGEMENTS

Thanks to Sivan Leviyang, Chris Illingworth, Uri Obolski, Kris Parag and Louis du Plessis for useful discussions.

## FUNDING STATEMENT

This work was supported by The Wellcome Trust and The Royal Society, grant number 107652/Z15/Z (RNT, JR and KL). RNT was supported further by Christ Church, University of Oxford, via a Junior Research Fellowship. CF and CW were supported by the Bridging the Evolution and Epidemiology of HIV in Europe (BEEHIVE) study via a European Research Council Advanced Grant, grant number PBDR-339251. The funders had no role in study design, data collection and analysis, decision to publish, or preparation of the manuscript.

## DATA AVAILABILITY

Computing code for running the model and data describing the distribution of variants in donors are accessible at https://github.com/robin-thompson/MultiplicityOfInfection

## COMPETING INTERESTS

The authors declare that no competing interests exist.

## GLOSSARY

*Set-point viral load (SPVL)*: The approximately stable viral load observed during chronic HIV-1 infection (see top left subpanel of Figure 1).
*Viral Load (HIV-1):*: The concentration of HIV-1 RNA in plasma, measured in copies per millilitre. We assume this is proportional to the concentration of viral particles in the blood.
*Viral Particle*:: An infectious unit, potentially consisting of either an individual virion or an infected cell.
*Viral Variant*:: A specified viral genotype (or group of genotypes). In this analysis, a viral variant refers to a unique genotype in the specific genomic region analysed.
*Virus:*: A single viral particle, or group of viral particles, all of the same variant. A new HIV-1 infection can be established by single or multiple viruses.

